# Complete plastid genome assembly of invasive plant, *Centaurea diffusa*

**DOI:** 10.1101/005900

**Authors:** Kathryn G. Turner, Christopher J. Grassa

## Abstract

New genomic tools are needed to elucidate the evolution of invasive, non-model organisms. Here we present the completed plastome assembly for the problematic invasive weed, *Centaurea diffusa*. This new tool represents a significant contribution to future studies of the ecological genomics of invasive plants, particularly this weedy genus, and studies of the Asteraceae in general.

## Introduction

Invasive species offer ample incentive and opportunity to address some of the major questions in evolution, ecology, and genetics. The direct costs of weed control and indirect costs of reduced crop production due to weeds are up to $40 billion annually in North America (Pimentel et al., 2005). Rates of adaptive phenotypic change are high in human-disturbed contexts (Hendry et al., 2008), such as invasion, and more common in introduced relative to native species in the same environment (Buswell et al., 2011). Thus, an ecological genomic approach may reveal key insights into a species’ adaptive capacity in the face of human-induced selective pressures.

This work aims to contribute to our genomic knowledge of the largest plant family, Asteraceae, and advance the study of invasion and evolution in *Centaurea diffusa*, a highly invasive weed species in North America. What genetic changes have occurred between native and invasive ranges of this species? Can these genetic changes be associated with adaptive trait shifts in the invaded range? To answer these questions, first several tools must be developed. Here we present one such tool, the complete plastid genome for *C. diffusa*, the first plastome from a genera containing approximately 250 species (Susanna and Garcia-Jacas, 2009), and one of only 10 genera in this speciose family with a complete plastome assembly.

## Methods

### Study Species

*Centaurea* species (knapweeds, star thistles) comprise the most abundant noxious weed genus in the western US, and are one of only 15 plant genera in the US significantly more likely to contain weedy species than expected by chance (Lejeune and Seastedt, 2001; Kuester et al., 2014). In the century since *Centaurea diffusa* was first reported there, it has formed dense monocultures, reduced forage quality, and altered soil and water resource availability in invaded grasslands (Lejeune and Seastedt, 2001). Recent work (Turner et al., 2014) has demonstrated the rapid evolution of *C.diffusa* in the invaded range under an array of benign and stressful conditions including drought. Invasive individuals grew larger, performed better, or matured later than native in nearly all tested conditions. Additionally, invasive individuals may have been released from a trade-off between growth and drought tolerance apparent in the native range.

### Collection and DNA extraction

Seed was collected from an individual in the native range of *C. diffusa* (TR001-1, Turkey, latitude 41.75111, longitude 27.24778). A voucher specimen of this population is located at the UBC Herbarium, accession number V236765. Seed from this collection was grown in a glasshouse at the University of British Columbia during the summer and fall of 2009. Seeds were germinated on filter paper in 1% plant preservative mixture and distilled H_2_O at room temperature. After 12 days seedlings were transplanted into 5 cm diameter cones filled with 80% potting mix and 20% silica sand. After two months, individuals were transplanted into 1 l pots containing potting soil to be used in a crossing experiment (“Maternal common garden”, Turner et al., 2014). Young leaf tissue was sampled from a single individual (TR001-1L) and stored at −80° C to be used for plastome assembly.

DNA was extracted from frozen tissue using a modified DNeasy column-less protocol. Concentration and quality was verified by Nanodrop, Qbit high-sensitivity assay, and gel electrophoresis.

### Plastome assembly and annotation

This whole genome shotgun library was sequenced at Genome Quebec, using one half lane of Illumina HiSeq 2000 paired-end sequencing. Raw data as well as scripts used for cleaning, assembly, and annotation of this plastome are available on Figshare.com (See Supplementary Materials). Raw reads were quality trimmed and screened for sequencing artifacts using Trimmomatic (Bolger et al., 2014). Clean reads were aligned to the *Lactuca sativa* plastome (Timme et al., 2007) using BWA (Li and Durbin, 2009). Pairs in which both reads aligned to the *L. sativa* plastome were extracted from the Sam files with Picard Tools SamToFastq.jar(Picard, 2009). ALLPATHS-LG (Gnerre et al., 2011) was used to merge overlapping pairs and error-correct the data, which was then assembled with Ray (Boisvert et al., 2010). Ray contigs were aligned to the *L. sativa* plastome with BLAST+ (Altschul et al., 1990) and scaffolded based on synteny using OSLay (Richter et al., 2007). Gaps were filled with GapFiller (Boetzer and Pirovano, 2012) resulting in a sequence containing a single N. Visual inspection indicated that the N separated an erroneous tandem duplication, which was corrected by hand with Vim. IR boundaries were confirmed with aTRAM (Allen and Huang, 2014) assemblies of flanking regions. Reads were aligned to the assembly with BWA and sorted with SAMTOOLS (Li et al., 2009). Visual inspection of the alignment revealed a few small indel and substitution errors, which were hand-corrected with Vim. A final alignment and inspection revealed no errors. The plastome was annotated using DOGMA (Wyman et al., 2004) and validated for NCBI GenBank submission using Sequin 13.05. The NCBI GenBank accession number for this plastome is KJ690264.

## Results

**Table 1:** Ray assembly output numerics

|  | Contigs | Contigs | Scaffolds | Scaffolds |
| --- | --- | --- | --- | --- |
| Size range | $\geq 100$ nt | $\geq 500$ nt | $\geq 100$ nt | $\geq 500$ nt |
| Number | 21 | 14 | 14 | 7 |
| Total length | 128037 | 126739 | 129501 | 128203 |
| Average | 6097 | 9052 | 9250 | 18314 |
| N50 | 23320 | 23320 | 27489 | 27489 |
| Median | 1234 | 2960 | 1112 | 11940 |
| Largest | 52890 | 52890 | 52890 | 52890 |

**Figure 1:**
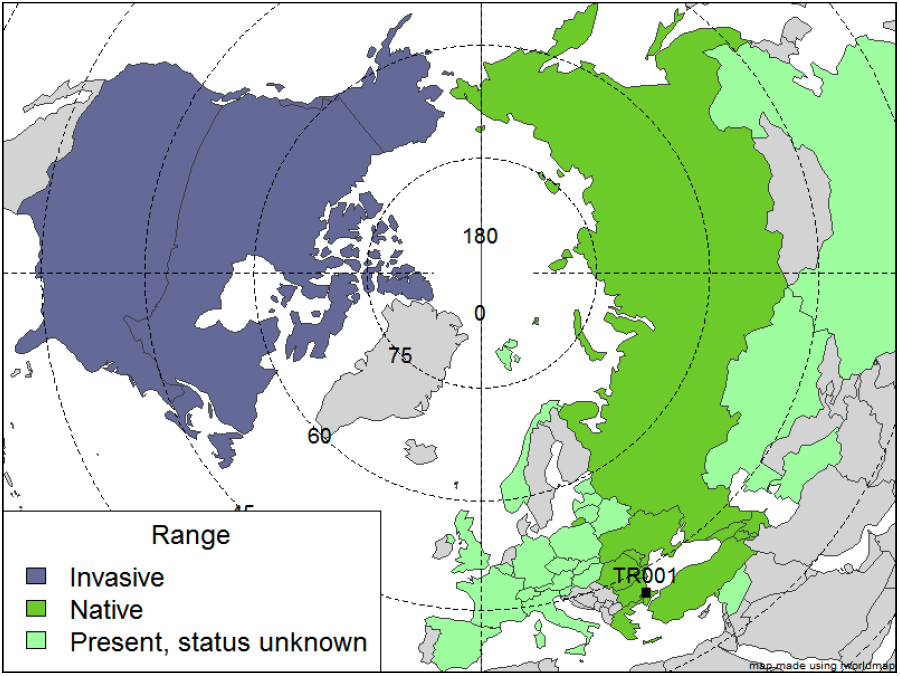
Global range map of *Centaurea diffusa*, by country (modified from Turner et al., 2014). Range status in a particular country is indicated by color. ‘Present, status unknown’ also includes countries where *Centaurea diffusa* is considered naturalized. Degrees of latitude are indicated on dotted lines, and degrees of longitude, solid lines. Seed collection location for the individual used in this assembly is indicated.

**Figure 2:**
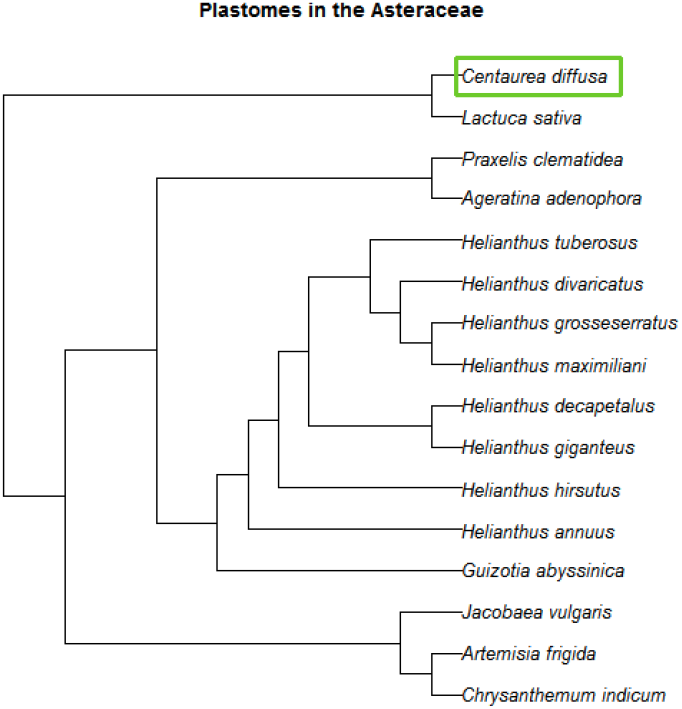
Phylogenetic tree of completed plastomes within the Asteraceae available on NCBI GenBank as of April 17, 2014. Phylogeny based on Smith et al. (2011), using R 3.0.1 (R core team, 2013), taxize (Chamberlain and Szocz, 2013), and Phylomatic (Webb and Donoghue, 2005).

**Figure 3:**
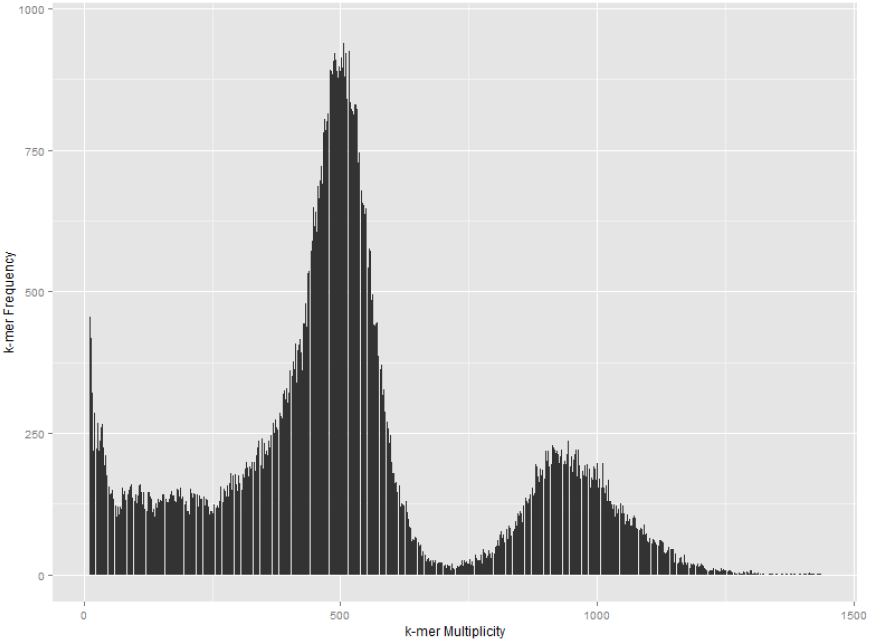
Histogram of 31-mers contained in *Centaurea diffusa* reads which aligned to *Lactuca sativa* plastome and their anchored orphans.

**Figure 4:**
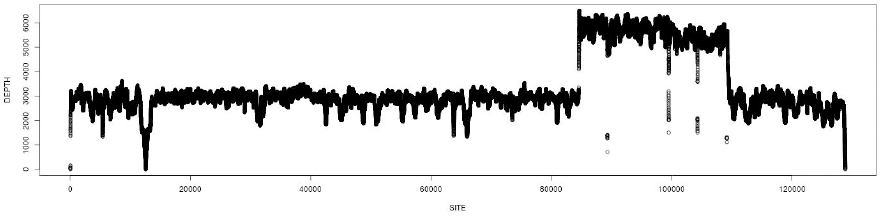
Depth of coverage of raw reads to *Centaurea diffusa* plastome v0.2 after gap filling which suggests: 1) misassembly near supercontig 1 position 12521, and 2) collapse of inverted repeat to a single copy.

**Figure 5:**
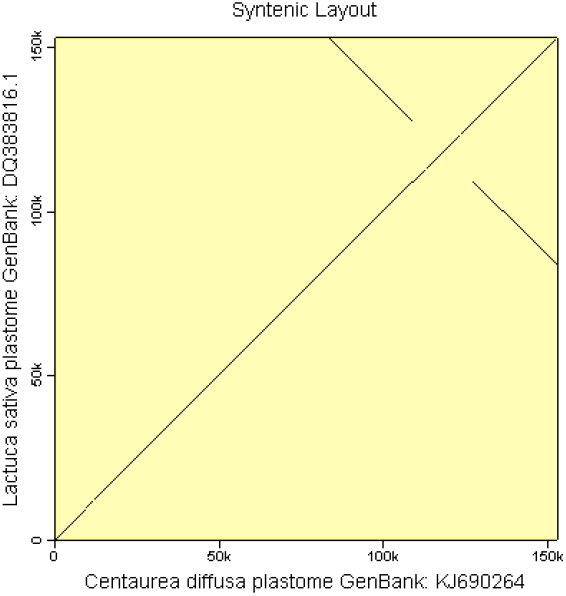
Syntenic alignment between *Lactuca sativa* and *Centaurea diffusa* plastomes using OSLay v1.0 (Richter et al., 2007).

**Figure 6:**
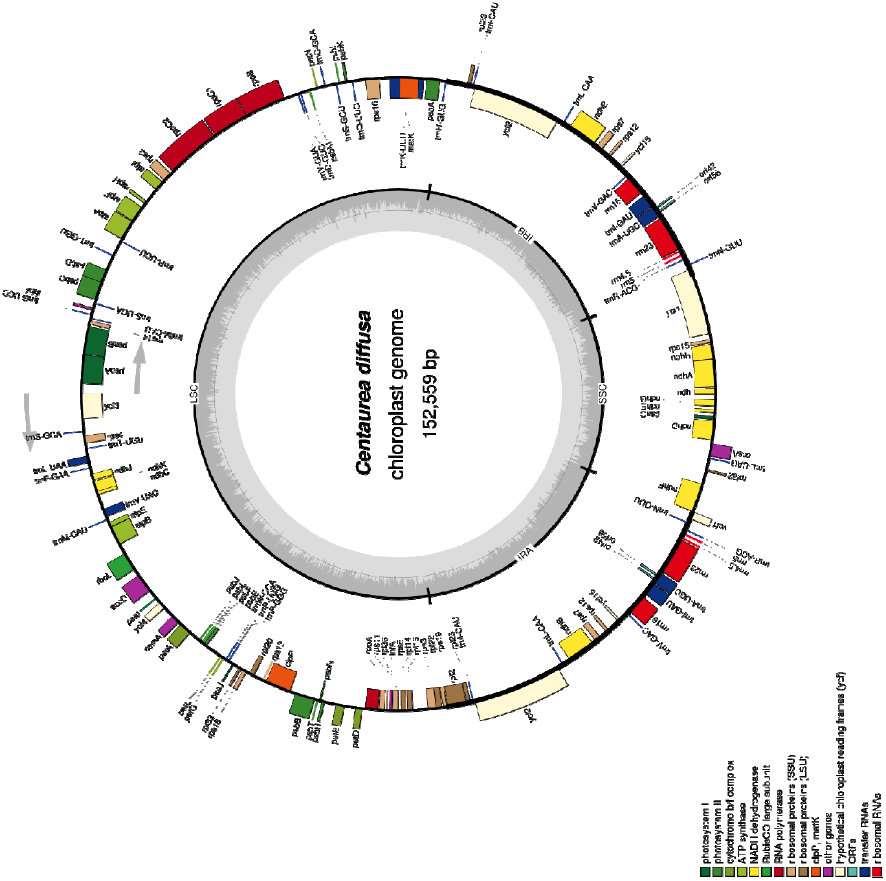
Map of annotated *Centaurea diffusa* plastome, produced using OGDraw (Lohse et al., 2013).

## Acknowledgements

We thank A. Guggisberg for seed collection, K. Nurkowski for plant care, D. Huang for assistance with error checking, and R. Timme for help with finalizing the annotation.

## Supplementary materials

Supplementary materials can be found at:

Complete plastid genome assembly of invasive plant, *Centaurea diffusa* - supplementary files. Kathryn Turner. fig**share**. Retrieved 22:17, Jun 03, 2014 (GMT) http://dx.doi.org/10.6084/m9.figshare.1044306

Supplementary files contain the following:

All figures.

Dataset S1: Raw reads that align to final assembly can be found in TR001_1L.to.TR001_1L.plastid.0.7.fa.sort.bam and TR001_1L.to.TR001_1L.plastid.0.7.fa.sort.bam.bai.

Scripts S1: All assembly code can be found in Centaurea_diffusa_Assembly_scripts.tar.gz. PLASTOME_MASTER.sh is the master script containing all commands and descriptions of manual steps.

Scripts S2: R code for the production of figures 1, 2, and 3.

